# Large dataset of octocoral mitochondrial genomes provides new insights into *mt-mutS* evolution and function

**DOI:** 10.1101/2021.09.08.459357

**Authors:** Viraj Muthye, Cameron D. Mackereth, James B. Stewart, Dennis V. Lavrov

## Abstract

All studied octocoral mitochondrial genomes contain a gene from the MutS family, whose members code for proteins involved in DNA mismatch repair, other types of DNA repair, meiotic recombination, and other functions. Although *mutS* homologues are found in all domains of life as well as viruses, octocoral *mt-mutS* is the only such gene encoded in an organellar genome. While the function of mtMutS is not known, its domain architecture, conserved sequence, and presence of some characteristic residues suggest its involvement in mitochondrial DNA repair. This inference is supported by exceptionally low rates of mt-sequence evolution observed in octocorals. Previous studies of *mt-mutS* have been limited by the small number of octocoral mt-genomes available. We utilized sequence-capture data from the recent Quattrini *et al*. study to assemble complete mitochondrial genomes for 97 species of octocorals. Combined with sequences publicly available in GenBank, this resulted in a dataset of 184 complete mitochondrial genomes, which we used to re-analyze the conservation and evolution of *mt-mutS*. We discovered the first case of *mt-mutS* loss among octocorals in one of the two *Pseudoanthomastus* sp. assembled from Quattrini *et al*. data. This species displayed accelerated rate and and changed patterns of nucleotide substitutions in mt-genome, which we argue provide additional evidence for the role of mtMutS in DNA repair. In addition, we found accelerated mt-sequence evolution in the presence of *mt-mutS* in several octocoral lineages. This accelerated evolution did not appear to be the result of relaxed selection pressure and did not entail changes in patterns of nucleotide substitutions. Overall, our results support previously reported patterns of conservation in *mt-mutS* and suggest that mtMutS is involved in DNA repair in octocoral mitochondria. They also indicate that the presence of *mt-mutS* contributes to, but does not fully explain, the low rates of sequence evolution in octocorals

## 1 Introduction

Over 1.5 billion years since their origin from *α*-proteobacteria, mitochondria have lost between 90% and 99% of their genes [1]. The remaining genes (just 37 in humans) [2] code for a handful of proteins – typically involved in the energy production – as well as a few structural RNAs necessary for the production of these proteins. While gene loss from the mitochondrial genome has been rampant, gene gain by mitochondria has been rare. In animals, such gain occurred only a handful of times, primarily in non-bilaterian animals (phyla Porifera, Cnidaria, Ctenophora, and Placozoa) (summarized in [3]). One of the best-known examples took place in octocorals, a group of marine cnidarians that includes soft corals, blue corals, sea pens, sea fans, and sea whips. All studied representatives of this group contain a mitochondrial gene (*mt-mutS*) coding for a MutS-like protein (mtMutS), a member of a large protein family involved in the DNA Mismatch Repair (MMR), other forms of DNA repair, and recombination [4, 5, 6].

The MutS protein family in animals is typically composed of five MutS homologues encoded in the nuclear genome (MSH2-6). Of these, three proteins – MSH2, MSH3, and MSH6 – function in nuclear MMR, while MSH4 and MSH5 function in meiotic recombination [7]. Recently, we found that an ortholog of yeast MSH1, a MutS protein targeted to mitochondria and involved in mitochondrial DNA repair [8, 9], is present in Hexacorallia, the sister group of octocorals, as well as two other phyla of non-bilaterian animals [10]. While the role of animal MSH1 has not been experimentally elucidated, its conserved structure, orthology to yeast MSH1, and inferred mitochondrial localization suggest a similar role in mitochondrial DNA repair [10].

Octocoral mtMutS is the only mitochondrial DNA(mtDNA)-encoded MutS homologue in eukaryotes. Comparative studies have shown that it is well conserved both in sequence and domain composition and retains amino-acid residues functionally important for mismatch identification [6, 12]. Furthermore, octocorals have some of the lowest rates of mitochondrial sequence evolution [3], an observation taken as an indirect support for the role of mtMutS in mitochondrial DNA repair. And yet the retention of *mt-mutS* over at least 470 million years of octocoral evolution is puzzling [13]. This is because DNA repair genes are usually among the first to be lost in resident genomes, *i*.*e*., genomes of obligate parasites, endosymbionts, and cellular organelles [14, 15]. In addition, little to no expression of *mt-mutS* was found in the study by Shimpi et al. [16]. Interestingly, both the notion of the low rate of mitochondrial sequence evolution in octocorals as well as that of the universal presence of *mt-mutS* in octocorals have been challenged recently [17, 18].

A recent study on Anthozoa evolution generated a large dataset of sequence capture data that included mitochondrial sequences [19]. We utilized this dataset to assemble mt-genomes for 97 species of octocorals. Combined with other sequences that became available on GenBank, this resulted in a nearly 6X increase in the number of available octocoral mt-genomes compared to the previous analysis performed in 2011 [6]. Here we used this dataset to re-evaluate the basic characteristics of *mt-mutS* and analyzed the conservation and evolution of this gene and the whole mt-genome across broad diversity of octocorals.

## 2 Results

### 2.1 Accelerated mitochondrial sequence evolution in multiple octocoral lineages

Phylogenetic analysis of concatenated amino acid sequences inferred from 184 complete octocoral mitochondrial genomes (mt-genomes) revealed eight species with highly accelerated rates of sequence evolution: *Calicogorgia granulosa, Cornularia pabloi, Erythropodium caribaeorum, Leptophyton benayahui, Melithaea erythraea, Muricella* sp., Pseudoanthomastus sp. 1 and *Tenerodus fallax* (Figure 2). Seven of these species were also represented by long branches on the phylogenetic tree based on mtMutS sequences (Figure S1), while one, Pseudoanthomastus sp. 1 lacked *mt-mutS* gene (see below). Although four of the fast-evolving mtMutS sequences grouped together on the MutS phylogenetic tree, this grouping was not present in the tree based on other coding sequences and likely represented the long branch attraction artifact [20]. Thus, acceleration in the rates of mitochondrial sequence evolution has occurred independently in multiple lineages of octocorals. In *Paragorgia* sp. USNM 1075769, an internal stop codon in *mt-mutS* resulted in a truncated protein of 340 amino acids. This species is not represented by a long branch in the phylogenetic tree based on the coding sequences of all mt-genes except *mt-mutS*.

**Figure 1:**
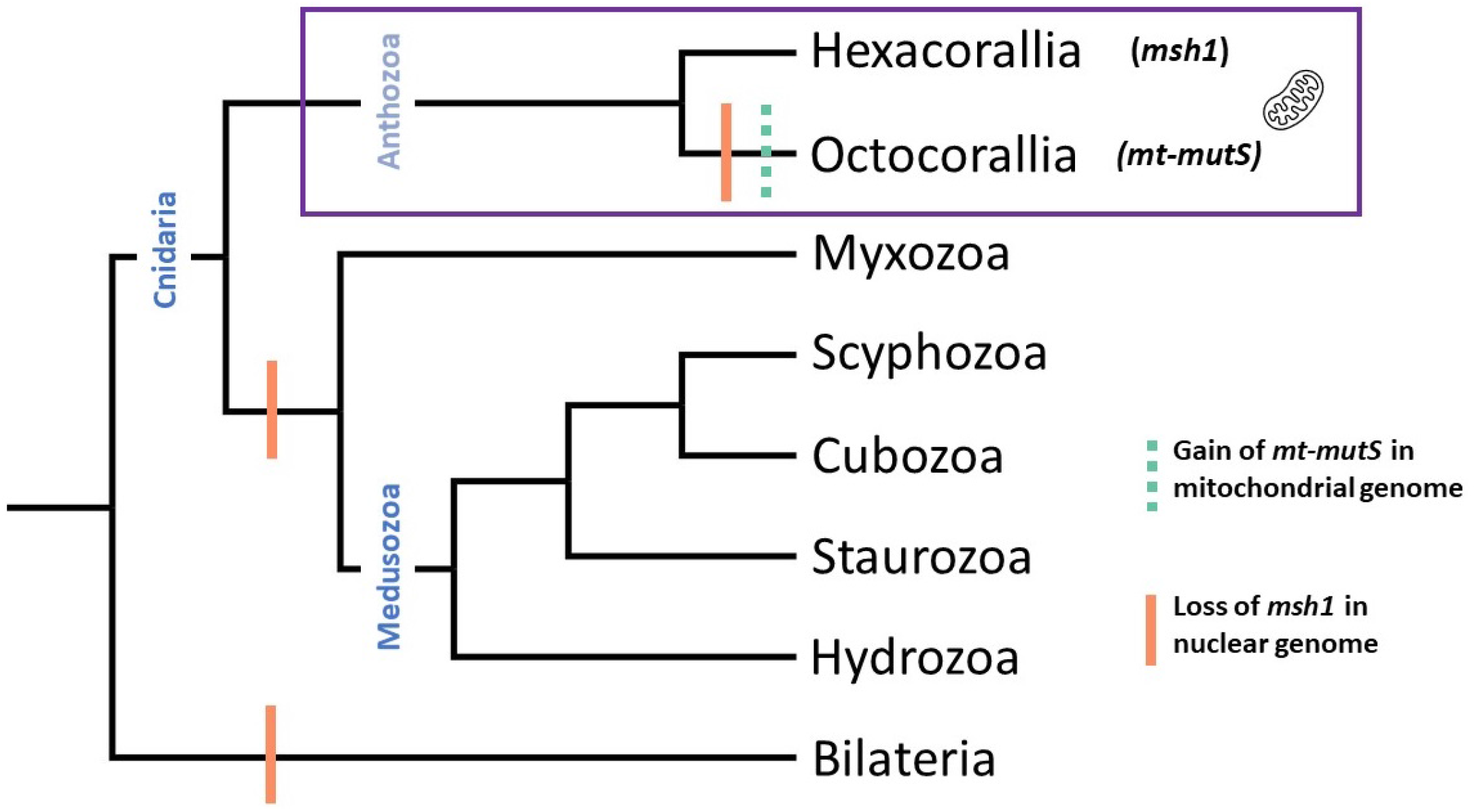
Loss of *msh1* and gain of *mt-mutS* in cnidarian groups. In the phylogenetic tree (adapted from [11]), the loss of *msh1* is shown by a orange bar and the gain of *mt-mutS* by a dashed green bar. The only cnidarian group that contains *msh1* (in Hexacorallia) and *mt-mutS* (in Octocorallia) – Anthozoa – is depicted by a purple box.

**Figure 2:**
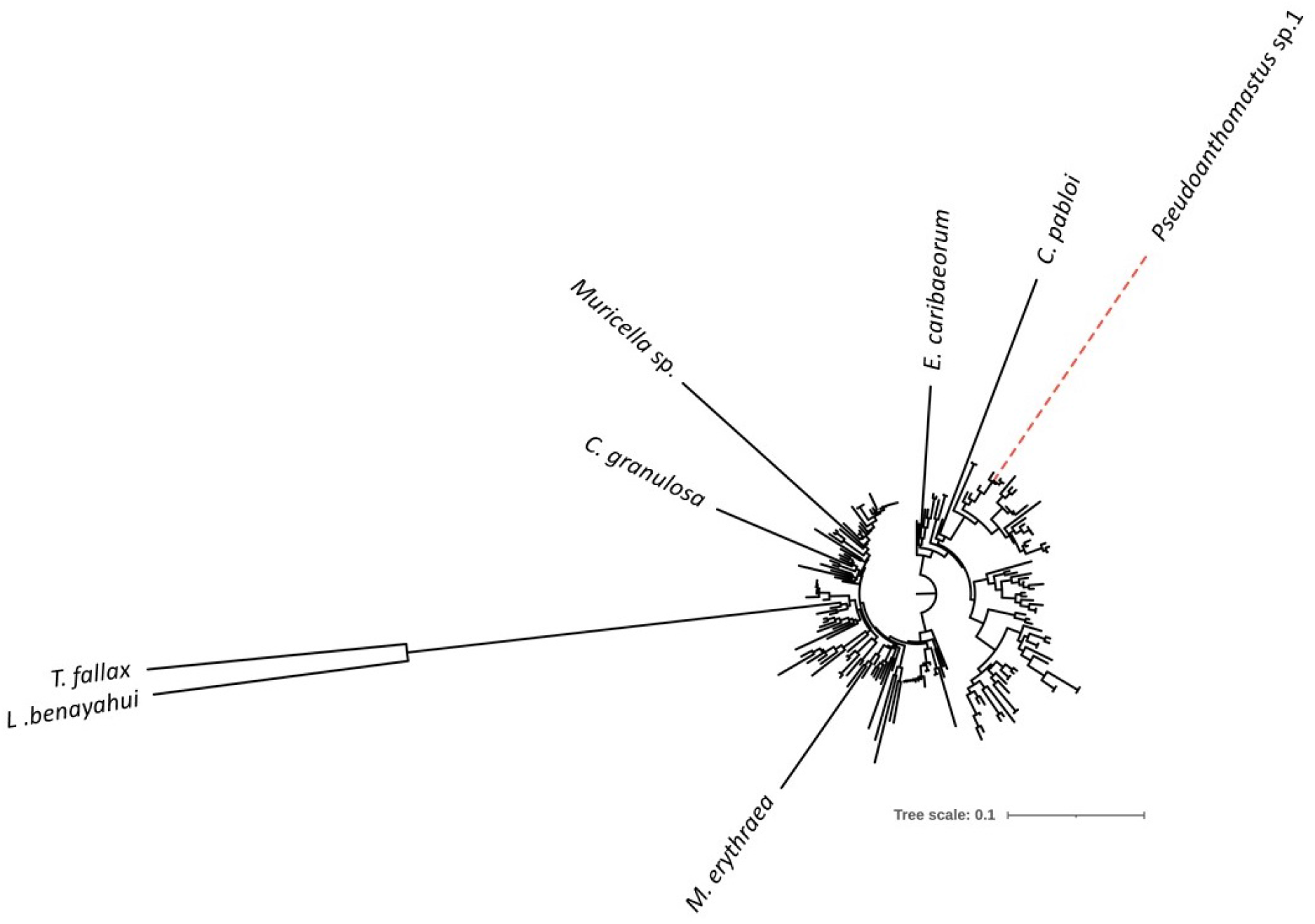
Maximum likelihood reconstruction of octocoral relationships based on concatenated amino acid sequences inferred from all mitochondrial protein genes except mtMutS. Named branches correspond to octocoral species with highly accelerated rate of mt-sequence evolution. Red dashed line indicates the only octocoral lacking *mt-mutS*.

### 2.2 Organization and conservation of octocoral mtMutS

Four protein domains were identified in all octocoral mtMutS sequences: “MutS I”, “MutS III”, “MutS V”, and “HNH”. In addition, the “MutS IV” domain, known to interact with DNA [21], was identified in 12 sequences, nested within the “MutS III” domain. All mtMutS sequences showed the same domain order: “MutS I - MutS III - MutS V - HNH”. The size of the protein domains ranged from 70 amino acids to 140 amino acids for “MutS I”, 297 amino acids to 376 amino acids for “MutS III”, 71 amino acids to 102 amino acids for “MutS IV”, 172 amino acids to 204 amino acids for “MutS V”, and 40 amino acids to 54 amino acids for “HNH”. The overall length of mtMutS proteins ranged from 957 amino acids in *Paragorgia* sp. 1075761 to 1,045 amino acids in *T. fallax* with the mean size of 988 amino acids. The GC content of *mt-mutS* ranged from 0.47 in *T. fallax* to 0.3 in multiple species. Species with the top four highest GC contenet were represented by long branches on the MutS phylogenetic tree – *T. fallax* (0.47), *L. benayahui* (0.45), *Muricella* sp. (0.42), and *C. pabloi* (0.39).

The sequence of the mtMutS proteins was well conserved among octocorals with the average pairwise sequence identity between any given species and the rest of octocorals ranging from 51.4% in *T. fallax* to 86.38% in *Eunicella tricoronata. L. benayahui* and *T. fallax*, two species with the largest distances also showed a large deviation in their amino acid composition (Figure 4A), being enriched in arginine and depleted in isoleucine and lysine (Table ST1). We identified 158 residues (15-16.5% of amino acids) conserved in all 182 mtMutS sequences. Among them was a phenylalanine residue at the N-terminus, critical for mismatch identification activity, as well as previously identified motifs inside ATPase and endonuclease domains. When excluding the seven fast evolving sequences, the number of 100% conserved positions increased to 242 (∼24% of the average-length protein). Within functional domains in mtMutS, the proportion of highly conserved residues (defined as those having a Consurf score of 7, 8, or 9) was 64.29%, 45.13%, 34.02%, 74.32%, and 52.08% for “MutS I”, “MutS III”, “MutS V”, and “HNH” respectively (Figure S4).

Nevertheless, mtMutS was one of the least conserved protein encoded by the octocoral mtgenome, with the second-lowest overall percentage of highly conserved residues (53.90%) in comparison to 51.56% (ATP8) - 84.95% (COX1) for others (Figure 3). Furthermore, mtMutS had the lowest GRAVY index value (a measure of hydrophobicity) among all mt-encoded octocoral proteins, ranging from -0.04 in *Paragorgia* sp. 1075761 to -0.28 in *Muricella* sp. (Figure S3).

**Figure 3:**
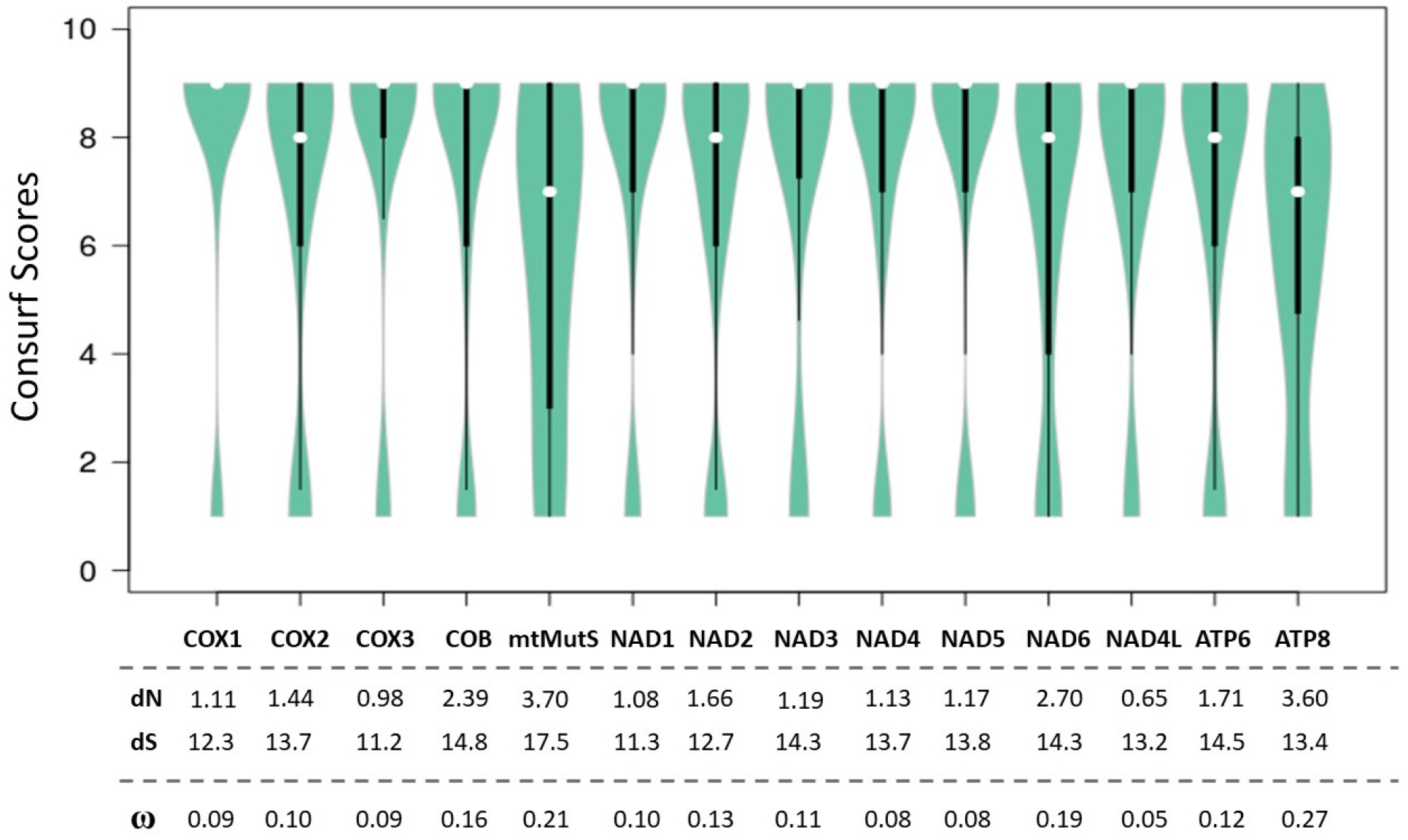
Distribution of Consurf scores in all fourteen mitochondrial proteins in octocorals. The number of positions in the alignment that have statistically significant scores are listed above each gene. The values for dN and dS for each gene calculated by CODEML are written below.

**Figure 4:**
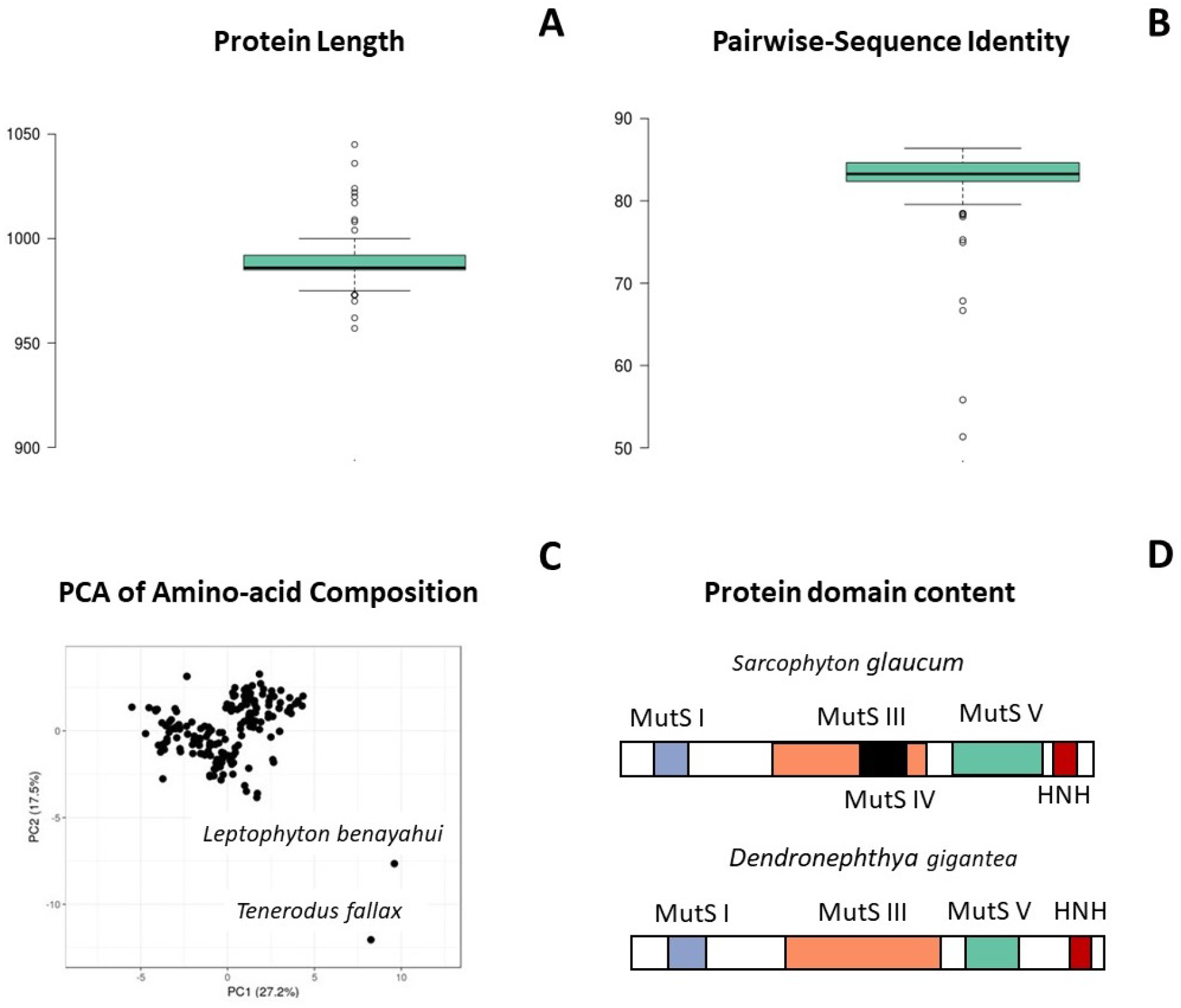
Basic characteristics of mtMutS in octocorals, A] Distribution of protein lengths of mtMutS, B] Distribution of GRAVY scores of mtMutS, C] Principle Component Analysis of the amino-acid composition of mtMutS, and D] Two types of protein domain architectures found in mtMutS proteins.

### 2.3 Rates of substitution and mode of selection

We calculated the rates of synonymous and non-synonymous substitutions for *mt-mutS* and compared them to those in other coding sequences in octocoral mt-genomes. While the rate of synonymous substitutions (dS) varied by 57% across the genes, ranging from 11.1 in *nad3* to 17.4 in *mt-mutS*, the rate of non-synonymous substitutions (dN) varied 6 times from 0.6 in *nad4L* to 3.66 in *mt-mutS*. The resulting *ω* values varied 4.6 times from 0.046 in *nad4L* to 0.21 in *mt-mutS* (Figure 3). The higher *ω* value in *mt-mutS* is consistent with lower conservation scores for mtMutS, indicating a relaxed purifying selection on more positions, but can also be influenced by positive selection acting on some codons. Indeed, by using site-specific models that allow *ω* to vary among sites, we identified seven sites under positive selection in *mt-mutS* (Table ST2). Four of these sites were in the “MutS III” domain; the remaining three were outside of functional domains. At each of these seven sites, identical amino-acid replacements have happened multiple times in octocorals. Some of these substitutions were conservative: at position 40, most sequences had proline or serine, which are both neutral, uncharged, and small amino-acids. However, at some positions, these replacements were radical: at position 33 there were multiple changes between isoleucine (nonpolar, uncharged, and relatively small) and lysine and arginine (both polar, relatively large, and positively charged.)

### 2.4 Increased rate and shifted pattern of substitutions in mitochondrial coding sequences of *Pseudoanthomastus* sp. 1

In contrast to other octocoral species analyzed in this study, no *mt-mutS* was found in the assembled mt-genome of *Pseudoanthomastus* sp. 1. Additionally, phylogenetic analysis based on other mitochondrial coding (mt-coding) sequences indicated that *Pseudoanthomastus* sp. 1 had a higher rate of mt-sequence evolution compared to the closely-related *Pseudoanthomastus* sp. 2, *Anthomastus* sp. USNM 1171062 and *Anthomastus* sp. USNM 1081145. The total estimated number of substitutions was ∼10X higher in *Pseudoanthomastus* sp. 1, compared to the other three species: 1516 vs. 137, 144, and 195 in *Pseudoanthomastus* sp. 2, *Anthomastus* sp. USNM 1171062, and *Anthomastus* sp. USNM 1081145, respectively. CODEML analysis found no significant change in the selection pressure on the branch leading to *Pseudoanthomastus* sp. 1 (p-value: 0.14), indicating a similar increase in both synonymous (mostly 3rd position) and non-synonymous (mostly 2nd and 1st positions) substitutions. By contrast, the increase in the number of transitions was much higher compared to the increase in transversions (Fig 5A), which resulted in an elevated *κ* value for *Pseudoanthomastus* sp. 1 (4.39), compared to *Pseudoanthomastus* sp. 2 (1.4), *Anthomastus* sp. USNM 1171062 (1.4) and *Anthomastus* sp. USNM 1081145 (1.12). As the result, 81% of all inferred substitutions in *Pseudoanthomastus* sp. 1 were transitions (Fig 5B). Interestingly, such a change has not been found in other fast-evolving species in the dataset. For example, a pairwise comparison between *T. fallax* and *L. benayahui* (Figure S5) revealed 1,500 transitions and 1,078 transversions, with a *κ* value of 1.39.

**Figure 5:**
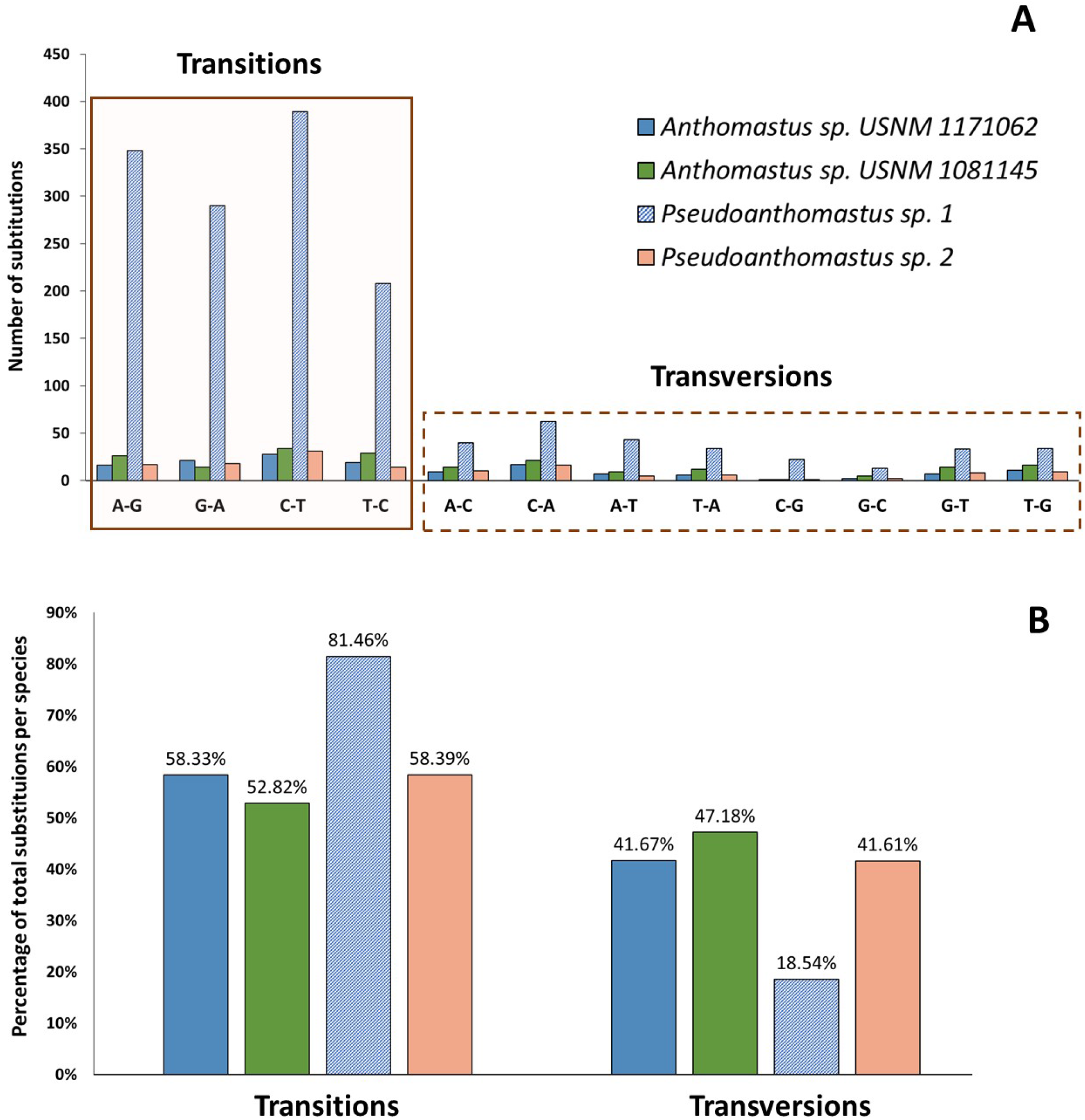
Analysis of patterns of substitution the mitochondrial genome of the only octocoral species without *mt-mutS* (*Pseudoanthomastus* sp. 1). This image shows individual substitution counts for each species of interest at all positions, compared to the outgroup species.

### 2.5 Sequence changes in mtMutS of octocoral species with accelerated rates of mt-genome sequence evolution

To understand whether any mtMutS changes might have contributed to the higher rates of mt-sequence evolution, we analyzed mtMutS sequences from the seven octocoral species that are represented by long branches in our phylogenetic trees by comparing them to the consensus of 175 slow(er)-evolving octocoral mt-MutS sequences. In particular, we analyzed the patterns of changes at 242 amino-acid positions that were conserved in these 175 mtMutS sequences. As reported above, 158 of these positions were universally conserved, while 84 had at least one amino acid replacement among fast evolving species. The number of changes ranged from seven in *M. erythraea* to 48 in *T. fallax*. The two species-*L. benayahui* and *T. fallax* had the most number of changes (37 and 48 respectively), of which 18 changes were found in both species. There were three positions (all within the “MutS III” protein domain) at which identical changes (deletions) occurred in all seven fast-evolving species (533L, 627L, and 628I in the consensus sequence).

## 3 Discussion

### 3.1 Is *mt-mutS* as well conserved as previously believed?

. The analysis of 30 octocoral mt-genomes by Bilewitch *et al*. [6] revealed several patterns of conservation in the octocoral mtMutS protein structure and sequence:

- all octocoral mt-genomes in that study contained *mt-mutS*
- all mtMutS proteins contained three domains: “MutS III” (the core domain), “MutS V” (the ATPase domain), and “HNH” (endonuclease domain) at the C-terminus; “MutS I”, the DNA mismatch recognition domain, was identified in all but three sequences
- several functionally important amino-acid position were found to be universally conserved

Our results based on a six times larger dataset generally support the results of Bilewitch et al. [6], with a few modifications. First and most importantly, we identified one species of octocorals – *Pseudoanthomastus* sp. 1 without *mt-mutS*, corroborating the unpublished study by [18]. Interestingly, this species belongs to a clade of octocorals with slow-evolving mtDNA, but experienced an accelerated rate of mt-sequence evolution in association with the *mt-mutS* loss. Second, we found the same domain composition of mtMutS as Bilewitch *et al*. [6], but provided evidence for the presence of an additional domain – “MutS IV” or the clamp domain – in 12 octocoral species. Interestingly, the “MutS IV” domain also shows an intermittent presence in other MutS proteins, like MSH1 [10]. Third, the functionally important amino-acids identified in [6] were well conserved in the expanded dataset, with a few exceptions. A phenylalanine residue (F44) necessary for mismatch recognition was found in 100% of the mtMutS as part of a “FYE” sequence. The “SVNGAGKST” motif of the ATPase domain (“MutS V”) was present in 172/182 mtMutS sequences. A glutamine residue (Q938) was universally conserved as part of the “LAQAG” sequence, as was the D1387 residue a part of the “LGDE” sequence, and H1425, a part of the “HLH” sequence. In the “HNH” endonuclease domain, H1202, H1203, and N1237 were 100% conserved, while H1246 was missing in just one sequence.

### 3.2 What is the function of mtMutS?

Bilewitch and Degnan [6] proposed that “the octocoral *mt-mutS* gene … codes for a protein with all the necessary components for DNA mismatch repair.” Indeed, the presence of the “MutS I” protein domain and the conserved phenylalanine residue required for mismatch identification in all mtMutS sequences, even from octocorals with fast-evolving mt-genomes, is striking. Across eukaryotic MutS subfamilies, the “MutS I” protein domain is found only in MutS homologues known to function in MMR: MSH2, MSH3, and MSH6, and is notably absent in MutS homologues that have non-MMR functions: MSH4 and MSH5. Furthermore, the universal presence of the HNH endonuclease domain within the protein likely makes it self-sufficient.

The tentative function of mtMutS in DNA repair is also supported by the change in the rate and patterns of substitutions in the mt-genome of *Pesudoanthomastus* sp. 1 (lacking *mt-mutS*) observed in this study: a large increase in transitions and only a slight increase in transversions. These patterns of substitution are consistent with the loss of some DNA repair capabilities as demonstrated by studies of mutation spectra in *E. coli* [22, 23, 24]. For example, in the study by [23], researchers carried out mutation accumulation analysis for several thousand generations of wild-type and MMR-defective *E. coli* (a strain that lacks the *mutL* gene). They saw that while transverions (that made up nearly 44% of mutations in the wild-type strain) increased 10-fold in MMR-deficient *E. coli*, transitions increased 200-fold. It is important to note that while a A:T *>* G:C bias was observed in the mutations that accumulated in MMR-deficient *E. coli*, a similar pattern was not observed in our analysis.

While the evidence discussed above supports the involvement of mtMutS in DNA repair, the type of the DNA repair is unkonwn. The main function of MMR is to remove sequence errors introduced during replication [25] and there is no evidence yet to link mtMutS activity to mtDNA replication. In fact, research on other systems argues against the involvement of mtMutS in MMR. Studies on yeast MSH1, a MutS homologue targeted to mitochondria have suggested that it is involved in BER instead of MMR (summarized in [26]). Another mitochondria-targeted MutS homologue – plant MSH1 – functions in a non-conventional DNA repair pathway. Plant MSH1 has a different endonuclease domain (“GIY-YIG”) at its C-terminus, which has been proposed to create a double-stranded break near the sites identified by “MutS I” protein domain, followed by homology-directed repair of the induced double-stranded break [27].

### 3.3 Should we be more surprised by the loss or retention of *mt-mutS*?

Our analysis revealed the presence of *mt-mutS* in 183 out of 184 analyzed octocoral mt-genomes. While the discovery of an octocoral species without *mt-mutS* was unexpected (but see [18]), we believe that one should be more surprised by the near universal retention of this gene across the octocoral diversity than spans more than 470 million years of evolution [13]. This is because genes involved in DNA repair are usually among the first to be lost in resident genomes [28]. An interesting example is the inactivation of the *recF* gene (involved in DNA recombinational repair) in the early stages of intracellular association in weevil endosymbionts [29]. Furthermore, the GRAVY scores (a measure of the hydrophobicity of the protein) of mtMutS are lower than those of all other mt-encoded proteins. This, according to the Hydrophobicity Hypothesis [30], should make it possible for *mt-mutS* to transfer to the nucleus and mtMutS to be imported by mitochondria. Although it is possible that the modified mt-genetic code in octocorals (UGA=Trp) provides an impediment to such transport, the UGA codon was only present in 72% of mt-mutS sequences (130/182) used for this study (see also [31]). Thus, it appears that there is a yet unknown functional constraint on mtMutS for its retention in octocoral mt-genomes.

### 3.4 Are mutation rates in *mt-mutS* higher than in other mt-genes?

Bilewitch and Degnan [6] also proposed that octocoral *mt-mutS* had a higher rate of mutations compared to other mitochondrial genes, in particular *cox1*, and found this observation paradoxical. However, their conclusion was based on pairwise sequence comparison for the two mitochondrial genes, which is not the right method for analyzing mutation rates. Pairwise distances (or sequence divergence) is the product of mutation rate, selection, and time and – when compared for different genes – are thought to reflect mainly differences in functional constraints on the encoded proteins. One way to disentangle the effects of mutation and selection is to analyze synonymous and non-synonymous substitutions separately, with synonymous substitutions used as a proxy for the mutation rate. This is because synonymous substitutions are often considered neutral and thus their rate should be equal to the mutation rate [32]. Indeed, our analysis showed that the rate of synonymous substitutions (dS) for *mt-mutS* and *cox1* are similar, while the average rate of non-synonymous substitutions (dN) is three time higher in *mt-mutS* compared to *cox1*. The higher average dS is likely a function of smaller percentage of highly conserved positions as indicated by its lowest conservation score among all mitochondrial genes. While multiple studies have reported that *mt-mutS* is transcribed [6, 16], the study by Shimpi *et al* [16] found that *mt-mutS* has very low expression levels. The low expression level of *mt-mutS* could explain the observation that it has the highest dS among all the octocoral mitochondrial genes.

### 3.5 Conclusions

In conclusion, our expanded sampling of octocoral *mt-mutS* showed that *mt-mutS* is 1) retained in all but one octocoral species, 2) well-conserved with respect to sequence and protein domain content, and 3) under purifying selection. mtMutS likely functions in DNA repair in octocoral mitochondria, but likely has an additional function remained to be characterized.

## 4 Methods and Materials

### 4.1 Assembling the datasets

We analyzed octocoral mitochondrial genomes from two datasets.

- **Dataset 1**: Mitochondrial genomes of 90 octocoral and 12 hexacoral species were downloaded from GenBank (Supplementary Materials File S1). The 12 hexacoral species were used as outgroups in our analysis. To ensure consistency in genome annotation across all species, we re-annotated all mitochondrial genomes using MITOS2 [33]. In case MITOS predicted multiple copies of the same gene in a genome, we retained the longest gene. In case a gene was not identified, we added the gene from the annotations in the GenBank files. The only gene missing from the assembled mitochondrial genomes is the *mt-mutS* gene from *Paragorgia* sp. USNM 1075769. An internal stop codon in this gene results in a truncated protein of 340 amino acids. Hence, we excluded it from all downstream analysis.
- **Dataset 2**: Mitochondrial genomes of 97 octocoral species were assembled from the data generated in [19]. The SRR files where downloaded with fastq-dump. Spades v. 3.14.0 was used for the sequence assembly [34]. For several species with high coverage raw data was sub-sampled with the “seqtk sample” command prior to assembly. Assembled mitochondrial contigs where identified by blast and extracted with the “seqtk subseq” command. The “mtann” script was used for mt-genome annotation.

We combined the *mt-mutS* gene and mtMutS protein sequences from the two datasets. We removed three sequences (*Complexum monodi, Corymbophyton bruuni*, and *Diodogorgia nodulifera*) since they had multiple partial genes.

### 4.2 Basic Characteristics of *mt-mutS*

We analyzed the following characteristics of the 182 *mt-mutS* sequences assembled:

- **Protein length**: Protein length distribution of mtMutS proteins was calculated using a custom script (“script1 find seqlength.bash”), and visualized using BoxPlotR [35].
- **Protein GRAVY score**: Protein GRAVY scores of mtMutS proteins were calculated using the GRAVY calculator at http://www.gravy-calculator.de/(last accessed: 08/26/2021 at 06:48PM) and visualized using BoxPlotR. The GRAVY score is a measure of the hydrophobicity of the protein, and is calculated as the sum of the hydropathy values of all amino acids divided by the length of the protein. GRAVY scores were calculated for all 14 mt-encoded octocoral proteins.
- **Amino-acid composition**: Protein amino-acid composition was calculated using a custom script (“script2 aacomposition.pl”), and PCA of the amino-acid composition was visualized using ClustVis [36].
- **Protein domain composition**: HMM models of the following domains were downloaded from the PFAM database – “MutS I”, “MutS II”, “MutS III”, “MutS IV”, “MutS V”, and “HNH”. HMMer v3.2.1 was used to scan the mtMutS sequences (e-value: 1e-03) for these protein domains.
- **Nucleotide composition**: Nucleotide composition of *mt-mutS* sequences was analyzed using the “compseq” program in EMBOSS, and the PCA of the amino-acid composition was visualized using ClustVis [36].
- **GC content**: The GC content of *mt-mutS* was calculated using the “geecee” program in EMBOSS and visualized using BoxPlotR.
- **Consensus sequence**: The “cons” package was used to build the consensus amino-acid sequence of mtMutS sequences.
- **Protein sequence similarity**: Custom script was used to calculate the average pairwise sequence identity for each 182 mtMutS sequences (“script4 percentage identity.bash”).

### 4.3 Phylogenetic analyses

We built phylogenetic trees of 1) complete mtMutS sequences (MutS phylogenetic tree) and 2) all mitochondrial proteins except mtMutS (Mitochondrial phylogenetic tree).

#### 4.3.1 MutS phylogenetic tree

The 182 complete mtMutS protein sequences were aligned using MAFFT v7.453 (using the –auto option) [37]. Poorly aligned regions in the alignment were trimmed using TrimAI (–automated1 option) [38]. The final alignment consisted of 182 sequences and 1,024 positions. This alignment is referred to as the “MutS dataset”. We used RAxML v8.2.11 [39] for the phylogenetic analysis of the final alignment. RAxML was used to build a phylogenetic tree for the resulting alignment with automatic selection of the substitution model and rapid bootstrapping with 1,000 resamples (“-m PROTGAMMAUTO -p 12345 -x 12345 -# 1000”). The resulting tree were visualized in iTOL v5.7 [40].

#### 4.3.2 Mitochondrial phylogenetic tree

We aligned all proteins (except mtMutS) from all 184 octocoral species using MAFFT (–auto option), and concatenated the individual alignments. We used TrimAI (-automated1 option) to remove poorly aligned regions. The final alignment comprised of 3,902 residues and is referred to as the “Mito dataset”. Like the MutS dataset, we used RAxML to build the phylogenetic tree for the Mito dataset. RAxML was used with automatic selection of the substitution model and rapid bootstrapping with 1,000 resamples (“-m PROTGAMMAUTO -p 12345 -x 12345 -# 1000”).

### 4.4 Conservation of octocoral *mt-mutS*

#### 4.4.1 Consurf analysis

Consurf was used to estimate position-specific conservation scores in octocoral mtMutS [41]. We used all the 182 mtMutS amino acid sequences in this analysis. The phylogenetic tree generated using RAxML was using for the Consurf analysis. These conservation scores generated by Consurf range from 1 (most variable) to 9 (most conserved). Consurf was also used to analyze all mitochondrial proteins in octocorals. We also scanned the complete mtMutS sequences for well-conserved functional residues summarized in [6].

#### 4.4.2 Analysis of patterns of selection in *mt-mutS* using CODEML

We aligned the protein sequences of all mitochondrial proteins using MAFFT (“–auto” option). The individual alignments were concatenated and then CD-HIT was used to remove sequences that were at least 97% similar. This resulted in a set of 89 octocoral mitochondrial genomes, and is hereby referred to as the “reduced dataset”.

CODEML was used to analyze the patterns of selection in mtMutS using the “reduced dataset”. MAFFT and Pal2Nal [42] were used to generate protein and codon alignments of octocoral mtMutS respectively. The codon alignment was used as input for CODEML [43] to analyze the signature of selection among codons in octocoral mtMutS. The ω(dn/ds) ratio is used as a measure of natural selection acting on a protein, with a ratio of 1, less than 1 and more than 1 indicating neutral evolution, purifying and positive selection respectively. Site models implemented in CODEML allow *ω* to vary among sites. Two Log-Likelihood ratio tests (LRTs) were carried out – M1 v/s M2a and M7 v/s M8 – to check for sites evolving under positive selection. Bayes Empirical Bayes (BEB) was used to identify sites under positive selection. Additionally, CODEML was used to calculate the *ω* value for all mitochondrial genes.

For examining patterns of substitutions, codon alignments of all protein-coding genes were used to identify nucleotides in a given species of interest to a chosen outgroup species, using a custom script (“script5 changes.py”). We compared the character states for each site, while tracking information on codon-position (first, second, or third). We examined whether the substitution was a transition or transversion. For each substitution, we also analyzed the direction of substitution (A-T or T-A).

#### 4.4.3 Data availibility

The scripts, sequences, and other data from this study were deposited in the repository at Open Science Framework at Muthye, V., Lavrov, D., 2021. Analysis of the octocoral mtMutS. doi:10.17605/OSF.IO/JZT8X.

## 6 Acknowledgement and Funding Sources

This study would not be possible without the sequence capture data generated by Quattrini *et al*. We thank Triston Walsh for his assistance with processing the data. Funding for this project was provided by a Research Grant (Ref.-No: RGP0050/2019) from the Human Frontiers Science Program.

## 7 Supplementary Figures

**Figure S1:**
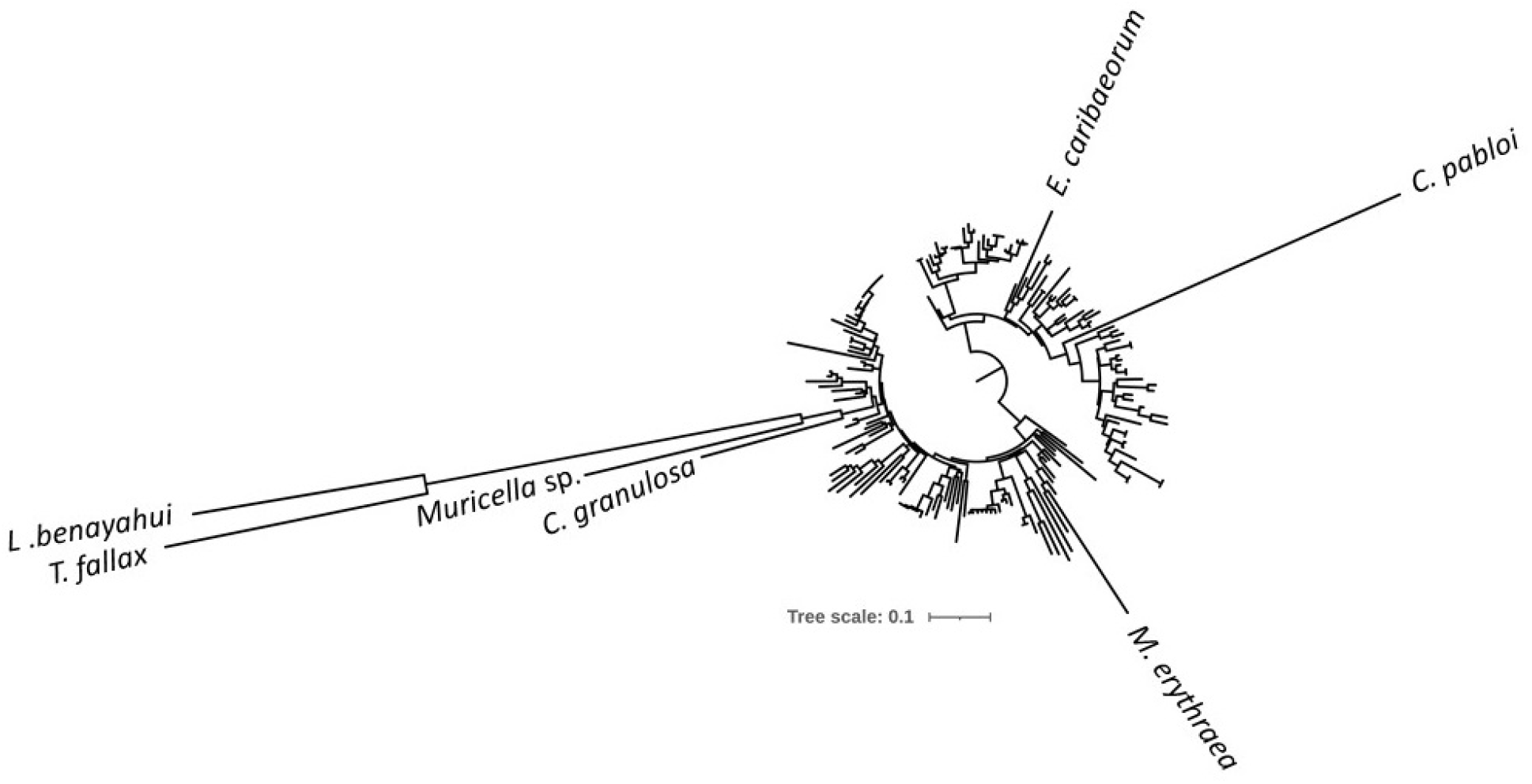
Phylogenetic tree of all octocoral mtMutS protein sequences generated using RAxML.

**Figure S2:**
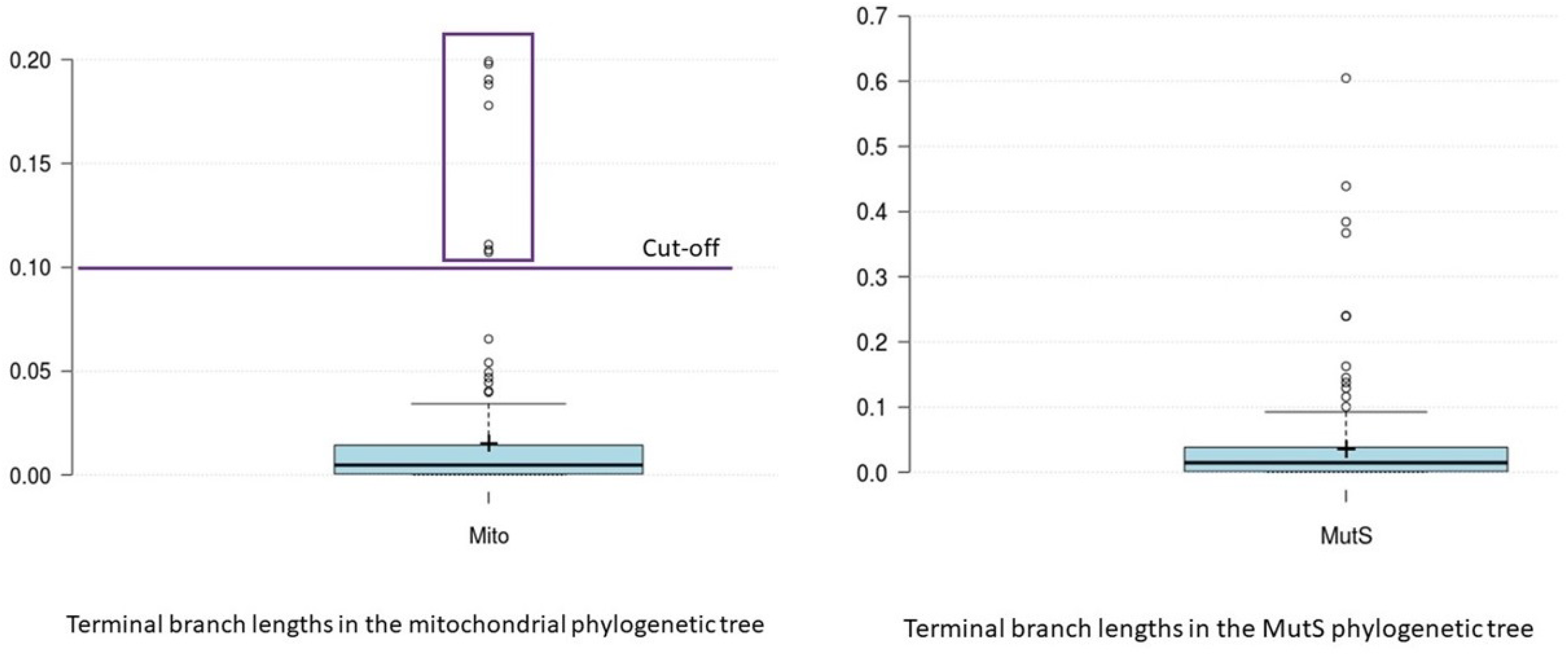
Distribution of terminal branch lengths in the mitochondrial and MutS phylogenetic tree.

**Figure S3:**
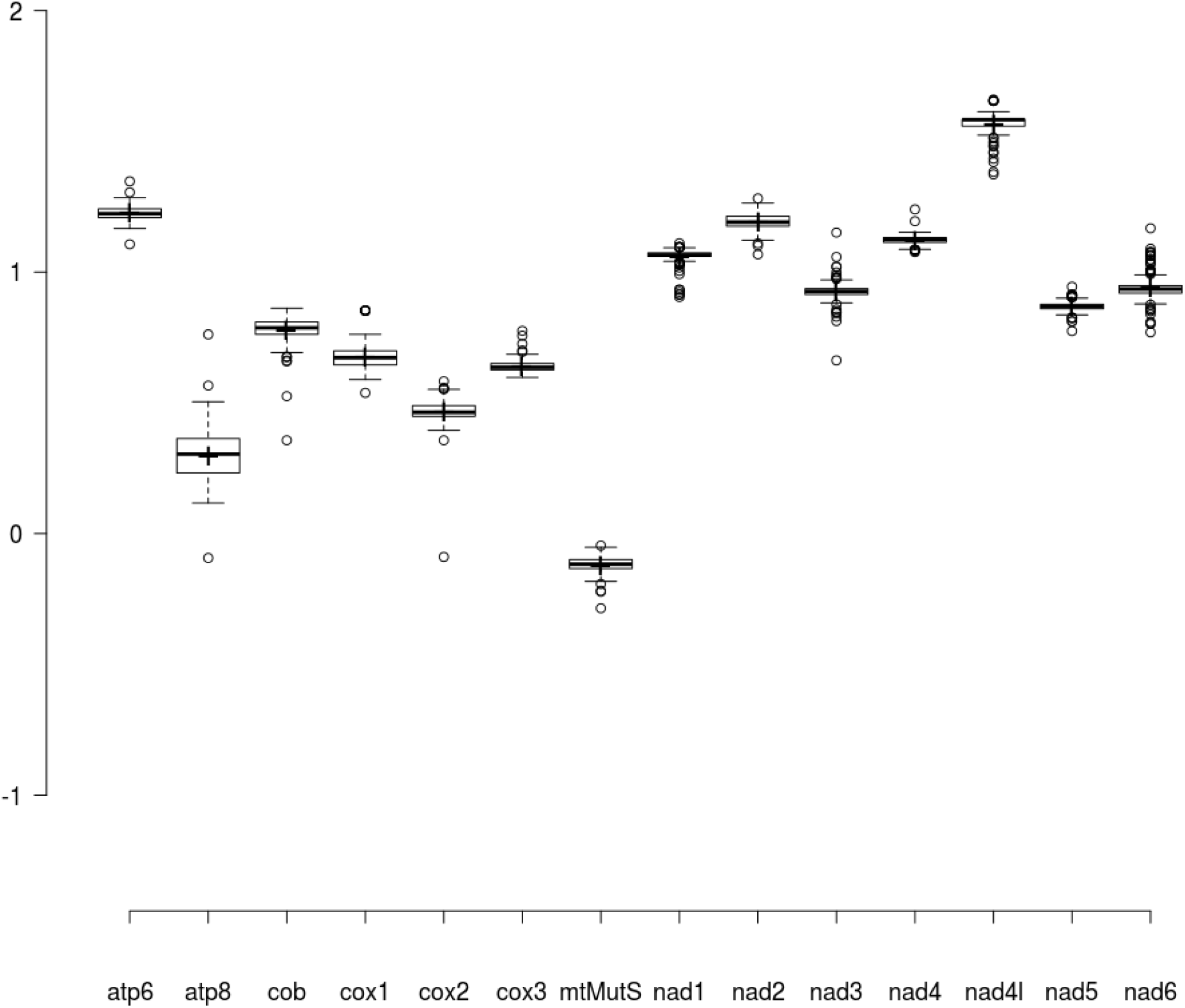
Distribution of the GRAVY index scores of mt-encoded proteins in octocorals.

**Figure S4:**
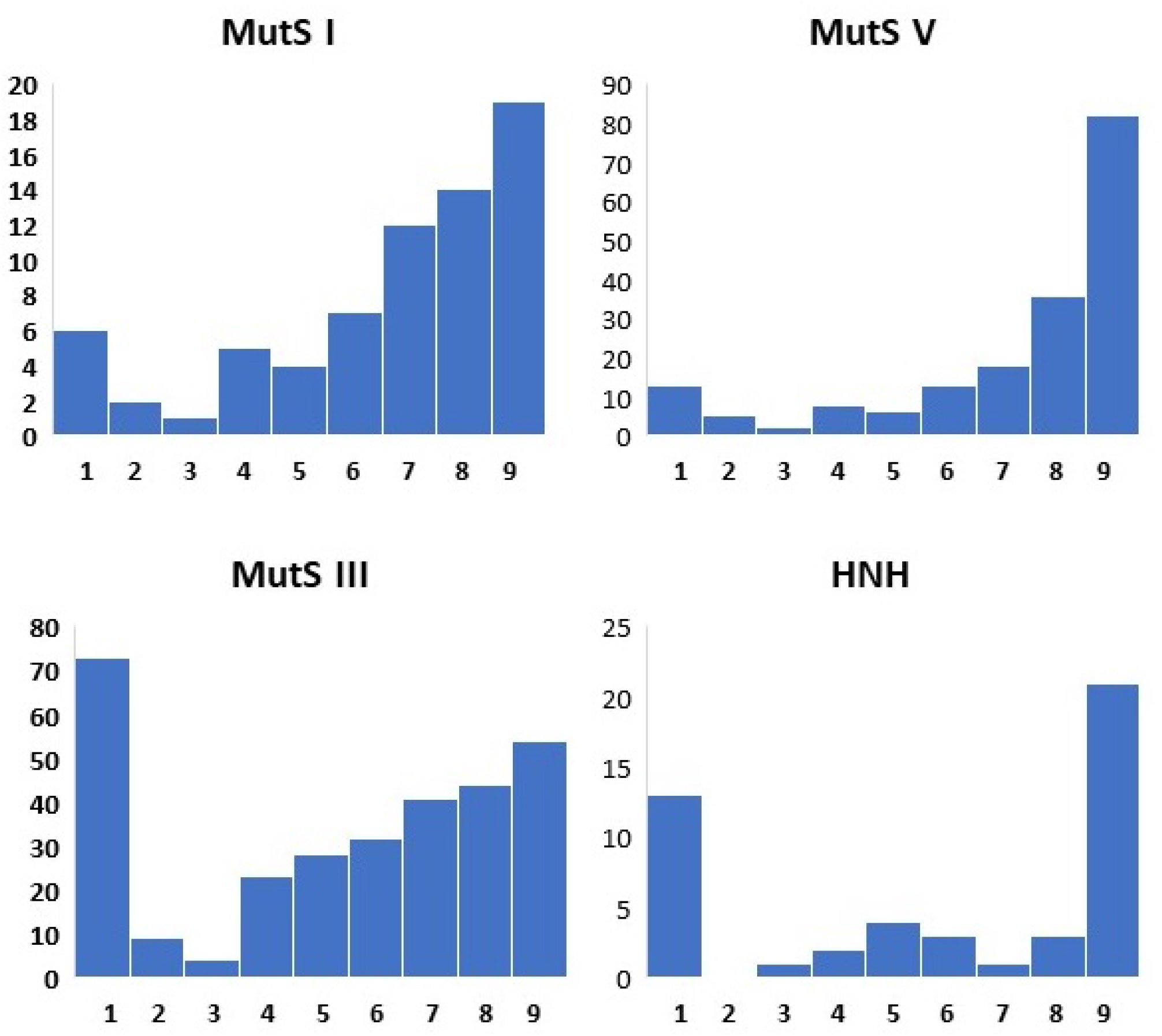
Consurf analysis of octocoral mtMutS. This is the distribution of the Consurf scores at each amino-acid residue within the four functional domains. The reference sequence used in this analysis was from *Sarcophyton glaucum*. The scores range from 1 (most variable) to 9 (least variable).

**Figure S5:**
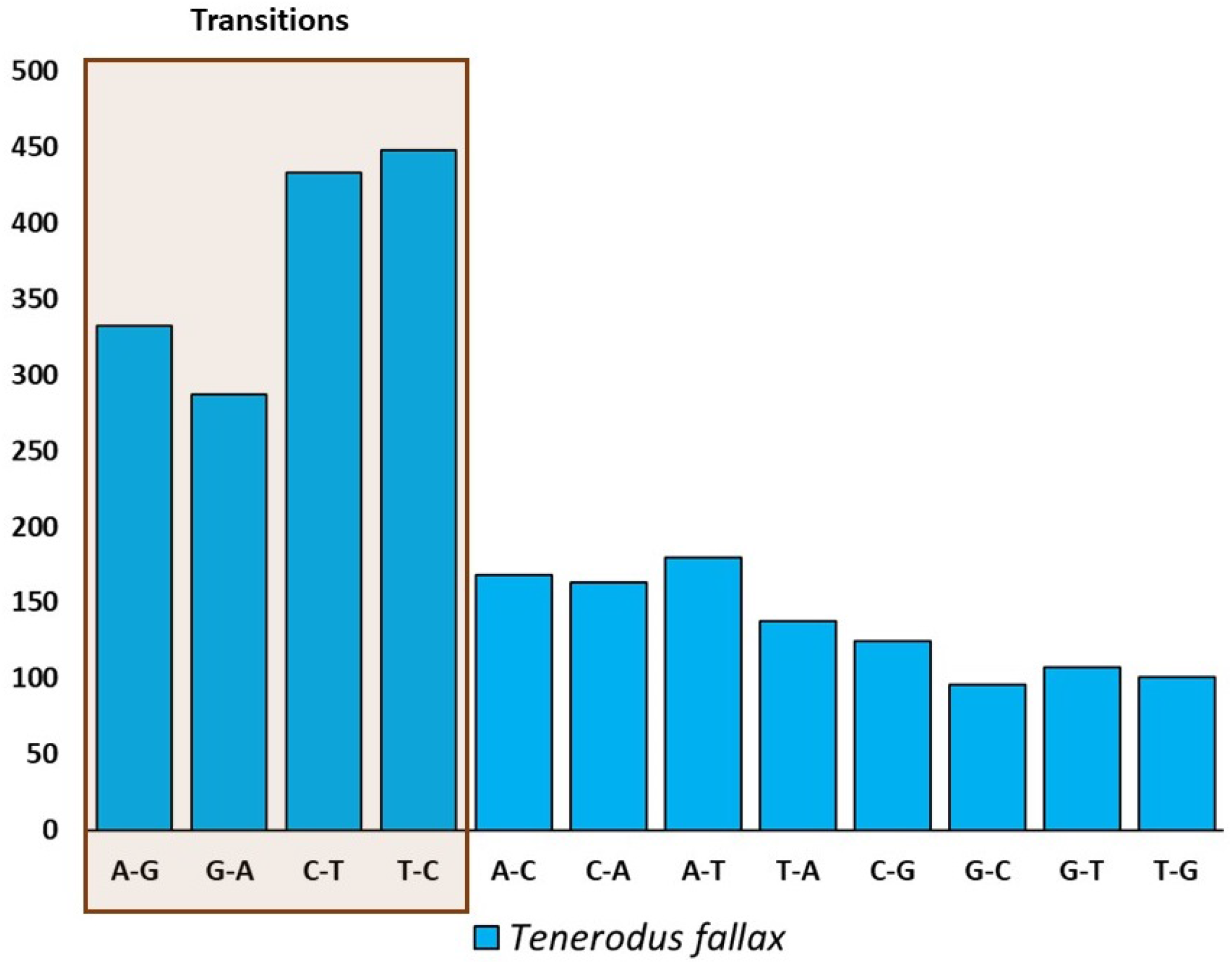
Analysis of patterns of substitution in *Tenerodus fallax* compared to *Leptophyton benayahui*

**Table ST1:** Comparison of the amino-acid content of mtMutS from *L. benayahui* and *T. fallax* to mtMutS from other octocoral species using Composition Profiler.

|  |  |  |
| --- | --- | --- |
| Ala | Not significant. | P-value=0.069544 (>0.001250) |
| Arg | Enriched. | P-value=0.000019 (≤0.001250) |
| Asn | Not significant. | P-value=0.003480 (>0.001250) |
| Asp | Not significant. | P-value=0.853725 (>0.001250) |
| Cys | Not significant. | P-value=0.641379 (>0.001250) |
| Gln | Not significant. | P-value=0.128119 (>0.001250) |
| Glu | Not significant. | P-value=0.278876 (>0.001250) |
| Gly | Not significant. | P-value=0.002550 (>0.001250) |
| His | Not significant. | P-value=0.008464 (>0.001250) |
| Ile | Depleted. | P-value=0.000008 (≤0.001250) |
| Leu | Not significant. | P-value=0.396715 (>0.001250) |
| Lys | Depleted. | P-value=0.000568 (≤0.001250) |
| Met | Not significant. | P-value=0.300812 (>0.001250) |
| Phe | Not significant. | P-value=0.326883 (>0.001250) |
| Pro | Not significant. | P-value=0.104908 (>0.001250) |
| Ser | Not significant. | P-value=0.231509 (>0.001250) |
| Thr | Not significant. | P-value=0.358851 (>0.001250) |
| Trp | Not significant. | P-value=0.525651 (>0.001250) |
| Tyr | Not significant. | P-value=0.013036 (>0.001250) |
| Val | Not significant. | P-value=0.001699 (>0.001250) |

**Table ST2:** Log likelihood ratio tests for the two model comparisons (M1 v/s M2a and M7 v/s M8) from CODEML.

| Model | lnL | LRT<br>Model comparison | Sites under positive<br>selection (BEB) |
| --- | --- | --- | --- |
| M1<br>(nearly neutral) | -45629.99 | p-value=5.72E-09 (df=2) | 33K*,40S,145T*,386E,387R*,419R,465H,511L,516M |
| M2A<br>(positive selection) | -45611.02 |  |  |
| M7<br>(beta) | -45156.32 | p-value=1.04E-31 (df=2) | 33K**,40S**,90V,113P,116P,145T**,166A,167N,210G,285R,<br>372Q,373L,383K,386E*,387R**,413E,419R,420A,465H**,471P,<br>472T,497Q,511L,516M**,538C,541K,753W,786L,787C |
| M8<br>(beta and<br>positive selection) | -45084.98 |  |  |

**Table ST3:**
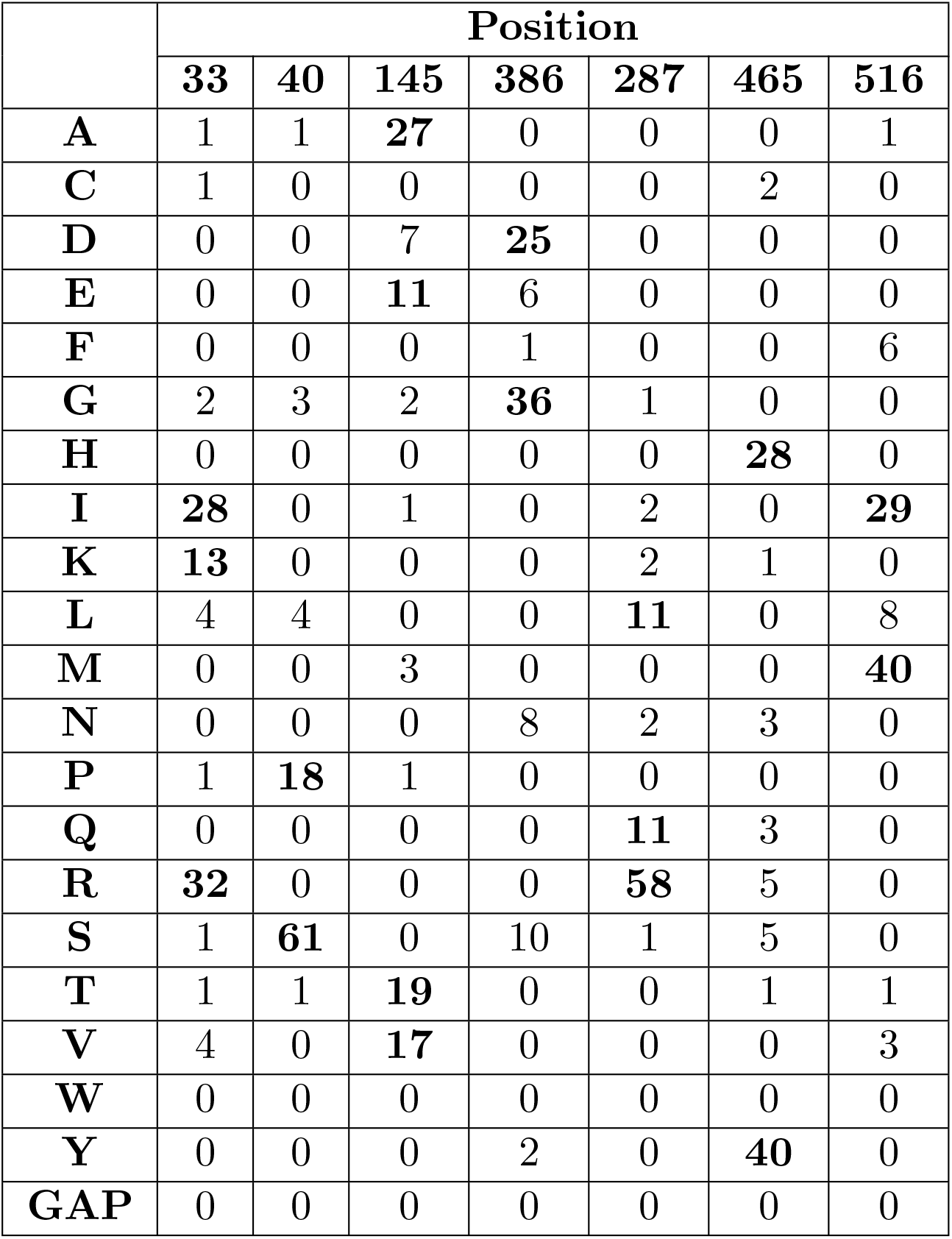
Distribution of amino acids at sites under positive selection in *mt-mutS*.

## References Cited

[1] Keith L Adams and Jeffrey D Palmer. Evolution of mitochondrial gene content: gene loss and transfer to the nucleus. Molecular phylogenetics and evolution, 29(3):380–395, 2003.

[2] Sharon Anderson, Alan T Bankier, Bart G Barrell, Maarten HL de Bruijn, Alan R Coulson, Jacques Drouin, Ian C Eperon, Donald P Nierlich, Bruce A Roe, Frederick Sanger, et al. Sequence and organization of the human mitochondrial genome. Nature, 290(5806):457–465, 1981.

[3] Dennis V Lavrov and Walker Pett. Animal mitochondrial dna as we do not know it: mt-genome organization and evolution in nonbilaterian lineages. Genome Biology and Evolution, 8(9):2896–2913, 2016.

[4] Geneviéve A Pont-Kingdon, Norichika A Okada, Jane L Macfarlane, C Timothy Beagley, David R Wolstenholme, Thomas Cavalier-Smith, and G Desmond Clark-Walker. A coral mitochondrial muts gene. Nature, 375(6527):109–111, 1995.

[5] Geneviáve Pont-Kingdon, Norichika A Okada, Jane L Macfarlane, C Timothy Beagley, Cristi D Watkins-Sims, Thomas Cavalier-Smith, G Desmond Clark-Walker, and David R Wolstenholme. Mitochondrial dna of the coral sarcophyton glaucum contains a gene for a homologue of bacterial muts: a possible case of gene transfer from the nucleus to the mitochondrion. Journal of molecular evolution, 46(4):419–431, 1998.

[6] Jaret P Bilewitch and Sandie M Degnan. A unique horizontal gene transfer event has provided the octocoral mitochondrial genome with an active mismatch repair gene that has potential for an unusual self-contained function. BMC evolutionary biology, 11(1):228, 2011.

[7] Jonathan A Eisen. A phylogenomic study of the muts family of proteins. Nucleic acids research, 26(18):4291–4300, 1998.

[8] RA Reenan and Richard D Kolodner. Characterization of insertion mutations in the saccharomyces cerevisiae msh1 and msh2 genes: evidence for separate mitochondrial and nuclear functions. Genetics, 132(4):975–985, 1992.

[9] P Koprowski, M Fikus, P Mieczkowski, and Z Ciesla. A dominant mitochondrial mutator phenotype of saccharomyces cerevisiae conferred by msh1 alleles altered in the sequence encoding the atp-binding domain. Molecular Genetics and Genomics, 266(6):988–994, 2002.

[10] Viraj Muthye and Dennis V. Lavrov. Dynamic evolution of the muts family in animals: multiple losses of msh paralogues and gain of a viral muts homologue in octocorals. bioRxiv, 2020.

[11] Ehsan Kayal, Bastian Bentlage, M Sabrina Pankey, Aki H Ohdera, Monica Medina, David C Plachetzki, Allen G Collins, and Joseph F Ryan. Phylogenomics provides a robust topology of the major cnidarian lineages and insights on the origins of key organismal traits. BMC evolutionary biology, 18(1):68, 2018.

[12] Hiroyuki Ogata, Jessica Ray, Kensuke Toyoda, Ruth-Anne Sandaa, Keizo Nagasaki, Gunnar Bratbak, and Jean-Michel Claverie. Two new subfamilies of dna mismatch repair proteins (muts) specifically abundant in the marine environment. The ISME journal, 5(7):1143–1151, 2011.

[13] John CW Cope. Octocorallian and hydroid fossils from the lower ordovician of wales. Palaeontology, 48(2):433–445, 2005.

[14] Nancy A Moran, John P McCutcheon, and Atsushi Nakabachi. Genomics and evolution of heritable bacterial symbionts. Annual review of genetics, 42:165–190, 2008.

[15] Siv GE Andersson and Charles G Kurland. Reductive evolution of resident genomes. Trends in microbiology, 6(7):263–268, 1998.

[16] Gaurav G Shimpi, Sergio Vargas, Angelo Poliseno, and Gert Wörheide. Mitochondrial rna processing in absence of trna punctuations in octocorals. BMC molecular biology, 18(1):1–16, 2017.

[17] Catherine S Mcfadden and LPV Ofwegen. Revisionary systematics of the endemic soft coral fauna (octocorallia: Alcyonacea: Alcyoniina) of the agulhas bioregion, south africa. Zootaxa, 4363(4):451–488, 2017.

[18] Rebecca Bentley. Mitochondrial genomes of cauliflower corals (nephtheidae), 2020.

[19] Andrea M Quattrini, Estefanía Rodŕguez, Brant C Faircloth, Peter F Cowman, Mercer R Brugler, Gabriela A Farfan, Michael E Hellberg, Marcelo V Kitahara, Cheryl L Morrison, David A Paz-García, et al. Palaeoclimate ocean conditions shaped the evolution of corals and their skeletons through deep time. Nature Ecology & Evolution, 4(11):1531–1538, 2020.

[20] Joseph Felsenstein. Cases in which parsimony or compatibility methods will be positively misleading. Systematic zoology, 27(4):401–410, 1978.

[21] Meindert H Lamers, Anastassis Perrakis, Jacqueline H Enzlin, Herrie HK Winterwerp, Niels de Wind, and Titia K Sixma. The crystal structure of dna mismatch repair protein muts binding to a g· t mismatch. Nature, 407(6805):711–717, 2000.

[22] Rajesh V Iyer, Shivranjani C Moharir, and Satish Kumar. Overexpression of muts impairs dna mismatch repair and causes cell division defect in e. coli. bioRxiv, 2020.

[23] Heewook Lee, Ellen Popodi, Haixu Tang, and Patricia L Foster. Rate and molecular spectrum of spontaneous mutations in the bacterium escherichia coli as determined by whole-genome sequencing. Proceedings of the National Academy of Sciences, 109(41):E2774–E2783, 2012.

[24] Roel M Schaaper and Ronnie L Dunn. Spectra of spontaneous mutations in escherichia coli strains defective in mismatch correction: the nature of in vivo dna replication errors. Proceedings of the National Academy of Sciences, 84(17):6220–6224, 1987.

[25] Oliver Fleck and Olaf Nielsen. DNA repair. Journal of Cell Science, 117(4):515–517, 02 2004.

[26] Jisha Chalissery, Deena Jalal, Zeina Al-Natour, and Ahmed H Hassan. Repair of oxidative dna damage in saccharomyces cerevisiae. DNA repair, 51:2–13, 2017.

[27] Zhiqiang Wu, Gus Waneka, Amanda K. Broz, Connor R. King, and Daniel B. Sloan. Msh1 is required for maintenance of the low mutation rates in plant mitochondrial and plastid genomes. Proceedings of the National Academy of Sciences, 117(28):16448–16455, 2020.

[28] John P McCutcheon, Bret M Boyd, and Colin Dale. The life of an insect endosymbiont from the cradle to the grave. Current Biology, 29(11):R485–R495, 2019.

[29] Colin Dale, Ben Wang, Nancy Moran, and Howard Ochman. Loss of dna recombinational repair enzymes in the initial stages of genome degeneration. Molecular biology and evolution, 20(8):1188–1194, 2003.

[30] Gunnar von Heijne. Why mitochondria need a genome. FEBS letters, 198(1):1–4, 1986.

[31] Walker Pett and Dennis V Lavrov. Cytonuclear interactions in the evolution of animal mitochondrial trna metabolism. Genome biology and evolution, 7(8):2089–2101, 2015.

[32] Motoo Kimura. The neutral theory of molecular evolution. Cambridge University Press, 1983.

[33] Matthias Bernt, Alexander Donath, Frank Jühling, Fabian Externbrink, Catherine Florentz, Guido Fritzsch, Joern Pütz, Martin Middendorf, and Peter F. Stadler. Mitos: Improved de novo metazoan mitochondrial genome annotation. Molecular Phylogenetics and Evolution, 69(2):313–319, 2013. Mitogenomics and Metazoan Evolution.

[34] Anton Bankevich, Sergey Nurk, Dmitry Antipov, Alexey A Gurevich, Mikhail Dvorkin, Alexander S Kulikov, Valery M Lesin, Sergey I Nikolenko, Son Pham, Andrey D Prjibelski, et al. Spades: a new genome assembly algorithm and its applications to single-cell sequencing. Journal of computational biology, 19(5):455–477, 2012.

[35] Michaela Spitzer, Jan Wildenhain, Juri Rappsilber, and Mike Tyers. Boxplotr: a web tool for generation of box plots. Nature methods, 11(2):121, 2014.

[36] Tauno Metsalu and Jaak Vilo. Clustvis: a web tool for visualizing clustering of multivariate data using principal component analysis and heatmap. Nucleic acids research, 43(W1):W566– W570, 2015.

[37] Kazutaka Katoh and Daron M Standley. Mafft multiple sequence alignment software version 7: improvements in performance and usability. Molecular biology and evolution, 30(4):772–780, 2013.

[38] Salvador Capella-Gutiérrez, José M Silla-Martínez, and Toni Gabaldón. trimal: a tool for automated alignment trimming in large-scale phylogenetic analyses. Bioinformatics, 25(15):1972– 1973, 2009.

[39] Alexandros Stamatakis. Raxml version 8: a tool for phylogenetic analysis and post-analysis of large phylogenies. Bioinformatics, 30(9):1312–1313, 2014.

[40] Ivica Letunic and Peer Bork. Interactive tree of life (itol) v4: recent updates and new developments. Nucleic acids research, 47(W1):W256–W259, 2019.

[41] Fabian Glaser, Tal Pupko, Inbal Paz, Rachel E Bell, Dalit Bechor-Shental, Eric Martz, and Nir Ben-Tal. Consurf: identification of functional regions in proteins by surface-mapping of phylogenetic information. Bioinformatics, 19(1):163–164, 2003.

[42] Mikita Suyama, David Torrents, and Peer Bork. Pal2nal: robust conversion of protein sequence alignments into the corresponding codon alignments. Nucleic acids research, 34(Suppl 2):W609–W612, 2006.

[43] Ziheng Yang et al. Paml: a program package for phylogenetic analysis by maximum likelihood. Computer applications in the biosciences, 13(5):555–556, 1997.

